# Tree-sequence recording in SLiM opens new horizons for forward-time simulation of whole genomes

**DOI:** 10.1101/407783

**Authors:** Benjamin C. Haller, Jared Galloway, Jerome Kelleher, Philipp W. Messer, Peter L. Ralph

**Affiliations:** Dept. of Biological Statistics and Computational Biology, Cornell University, Ithaca, NY 14853, USA; Institute of Ecology and Evolution, University of Oregon, Eugene, OR 97403, USA; Big Data Institute, Li Ka Shing Centre for Health Information and Discovery, University of Oxford, Oxford, OX3 7FZ, UK

**Keywords:** pedigree recording, coalescent, background selection, genealogical history, selective sweeps, tree sequences

## Abstract

There is an increasing demand for evolutionary models to incorporate relatively realistic dynamics, ranging from selection at many genomic sites to complex demography, population structure, and ecological interactions. Such models can generally be implemented as individual-based forward simulations, but the large computational overhead of these models often makes simulation of whole chromosome sequences in large populations infeasible. This situation presents an important obstacle to the field that requires conceptual advances to overcome. The recently developed tree-sequence recording method (Kelleher et al., 2018), which stores the genealogical history of all genomes in the simulated population, could provide such an advance. This method has several benefits: (1) it allows neutral mutations to be omitted entirely from forward-time simulations and added later, thereby dramatically improving computational efficiency; (2) it allows neutral burn-in to be constructed extremely efficiently after the fact, using “recapitation”; (3) it allows direct examination and analysis of the genealogical trees along the genome; and (4) it provides a compact representation of a population’s genealogy that can be analyzed in Python using the msprime package. We have implemented the tree-sequence recording method in SLiM 3 (a free, open-source evolutionary simulation software package) and extended it to allow the recording of non-neutral mutations, greatly broadening the utility of this method. To demonstrate the versatility and performance of this approach, we showcase several practical applications that would have been beyond the reach of previously existing methods, opening up new horizons for the modeling and exploration of evolutionary processes.

## Introduction

Forward simulations are increasingly important in population genetics and evolutionary biology. For example, they can be useful for modeling the expected evolutionary dynamics of real-world systems (Fournier-Level et al., 2016; Cotto et al., 2017; Matz et al., 2018; Ryan et al., 2018), for discovering the ecological and evolutionary mechanisms that led to present-day genomic patterns in a species (Enard et al., 2014; Nowak et al., 2014; Arunkumar et al., 2015; Patel et al., 2018), for testing or validating empirical and statistical methods (Haller and Hendry, 2013; Caballero et al, 2015; Ewing et al., 2016; Haller and Messer, 2017a), and for exploring theoretical ideas about evolution (Haller et al., 2013; Assaf et al., 2015; Mafessoni and Lachmann, 2015; Champer et. al, 2018), among other purposes. Because of this broad utility, there is a growing desire to run simulations with increased realism in a variety of areas: longer genomic regions up to the scale of full genome sequences, large populations, selection at multiple loci with linkage effects, complex demography, ecological interactions with other organisms and the environment, explicit space with continuous landscapes, spatial variation in environmental variables, spatial interactions such as competition and mate choice between organisms, and so forth.

However, this type of realism comes at a price, in both processing time and memory usage. Since computational resources are finite, this can often make it difficult or, in practical terms, impossible to run some models. Advances in computing power have gradually extended the boundaries of what is possible, as have performance improvements due to improved forward simulation software (Messer, 2013; Thornton, 2014; Haller and Messer, 2017b), but computational overhead continues to hold back progress in the field by limiting the level of realism that can be attained in models.

From this perspective, the recently developed pedigree recording or “tree-sequence recording” method (Kelleher et al., 2018) is potentially transformative. Kelleher et al. (2018) have shown that, perhaps counterintuitively, the recording of all ancestry information for the entire population can actually improve the runtime by orders of magnitude. These gains in efficiency are made possible by the succinct tree sequence data structure (or “tree sequence”, for brevity) that lies at the heart of the msprime coalescent simulator (Kelleher et al., 2016), subsequently refined in Kelleher et al. (2018). The tree sequence data structure is a concise encoding of the correlated genealogies along a chromosome resulting from evolution in sexually reproducing populations (Figure 1). The sequence of trees along a genome has been studied for some time (Hudson, 1983), and is closely linked to the concept of an “Ancestral Recombination Graph” or ARG (Griffiths, 1991; Griffiths & Marjoram, 1997). The use of the term “ARG” has historically been ambiguous, however, sometimes referring to the stochastic process generating these trees, rather than the resulting tree sequence itself, so we use the term “tree sequence” here to refer to this sequence of trees in the particular representation described by Kelleher et al. (2016, 2018). Precisely the same tree sequence data structure can be used to record each generation’s parent–child relationships. This data structure will then record who each individual inherited each section of chromosome from, for every individual that ever lived. However, there is a massive amount of redundancy in this information, since many of the individuals simulated in the past will leave no descendants in the extant population. The key insight of Kelleher et al. (2018) was to provide an efficient algorithm to remove this redundancy by periodically “simplifying” the tree sequence. This combination – the tree sequence data structure and an efficient algorithm for simplifying it – allows complete genealogies for all extant individuals to be recorded efficiently in forward simulations for the first time.

**Figure 1.**
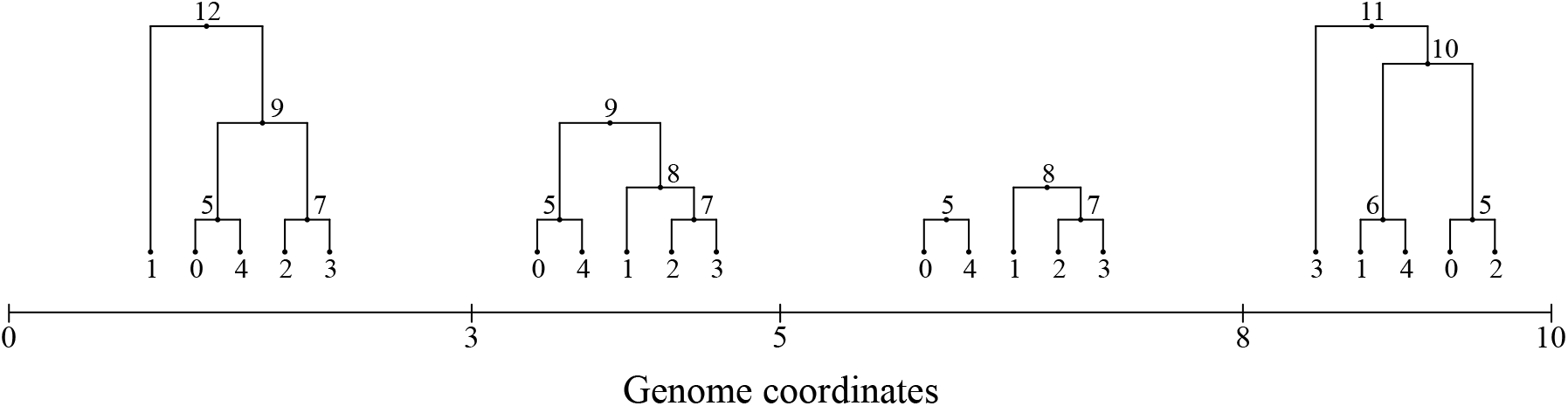
An example tree sequence for a model of five extant genomes, with a chromosome ten base positions long. Each interval between *x* axis ticks is a genomic interval with a distinct ancestry tree. The leaves of each tree [0–4] represent the extant genomes, whereas the internal nodes [5–12] represent ancestral genomes from which the extant genomes descend. The pattern of ancestry at adjacent sites is typically highly correlated, as seen here. Full coalescence has been achieved for the first, second, and fourth intervals, but the third interval has not yet fully coalesced; the tree for that interval therefore has multiple roots. See Kelleher et al. (2016, 2018) for further discussion of the tree sequence data structure.

The most immediate advantage of recording a tree sequence during forward simulation is that it allows neutral mutations to be omitted entirely; neutral mutations can simply be overlaid onto the tree sequence after forward simulation has completed, because by definition they do not affect the genealogies. This provides an immense efficiency benefit, since neutral mutations then only need to be added along those branches of the tree from which the individuals of interest at the end of the simulation have inherited; all other ancestral branches, which typically comprise the vast majority of the full tree, can be ignored since they do not contribute to those individuals. Given that many forward simulations spend the large majority of their time managing neutral mutations, with considerable bookkeeping overhead in each generation, neutral mutation overlay following forward simulation has been shown to improve performance by an order of magnitude or more while producing provably statistically identical results (Kelleher et al., 2018).

A second advantage of recording genealogies is that the recorded tree sequence from a forward simulation can be used as the basis for the construction of a neutral “burn-in” history for the simulated population after forward simulation is complete, using (usually much faster) coalescent simulation. The burn-in period of a simulation can be immensely time-consuming, often taking much longer than the simulation of the evolutionary dynamics that are actually of interest; the overhead of burn-in can therefore present a large obstacle for many models. With a method that we call “recapitation”, we can leverage the information in the tree sequence to prepend a coalescent simulation of the burn-in period, speeding up the burn-in process by many orders of magnitude.

A third important advantage is that the pattern of ancestry and inheritance is in itself very useful. For many statistics of interest, and in particular for inferring specific events that occurred in the past, sequence-based data from mutations is essentially an extra layer of noise over the signal of interest contained in the genealogies. Direct access to the precise genealogical history of the simulated population allows the signal to be analyzed without the noise, gaining significant statistical power. An expanding set of open-source tools makes it possible to load, analyze, and even manipulate a recorded tree sequence using simple Python code, allowing open-ended flexibility in analysis.

A fourth compelling advantage is that the recorded tree sequence files are very small and enable very efficient calculation of population-genetic statistics (Kelleher et al. 2016, 2018). The files output from even the largest simulations are rarely bigger than a few hundred megabytes, and may be tens of thousands of times smaller than alternatives such as VCF and Newick. Despite this high level of compression, tree sequences can be processed very efficiently; statistics of interest such as allele frequencies within cohorts can often be computed incrementally, leading to very efficient algorithms (Kelleher et al. 2016). Calculation of statistics of this sort from simulated data can be very time-consuming, especially when long genomes are involved and many replicate simulation runs have been performed, so the ability to speed up such calculations is quite important.

Given these advantages, we have worked to integrate tree-sequence recording into SLiM 3, a new major release of the free, open-source SLiM simulation software package (http://messerlab.org/slim/). It is now possible to enable tree-sequence recording in any SLiM model with a simple flag set in the model’s script, and then to output the recorded tree sequence at any point in the simulation. In addition, we have extended the original tree-sequence recording method (Kelleher et al. 2018) to allow for the recording of mutations during forward simulation. This allows the tree-sequence output format, a `.trees` file, to be used in SLiM as a way of saving and then restoring the state of a simulation while preserving information about ancestry, and allows the mutations that occurred during forward simulation to be accessed later in Python-based analyses.

To illustrate the large advantages provided by tree-sequence recording, and to show how to take advantage of those benefits when using SLiM for forward simulation, we will present four practical examples of the method. In the first example, we will show the impressive performance benefits that can be achieved with tree-sequence recording compared to a classical forward simulation. The second example will use tree-sequence recording to efficiently simulate background selection near genes undergoing deleterious mutations, quantifying the expected effect of background selection on levels of neutral diversity by measuring the heights of trees in the recorded tree sequence. Our third example will be a model of admixture between two subpopulations, showing how to use the recorded tree sequence in calculating the mean true local ancestry at every position along a chromosome. Finally, the fourth example will illustrate how the “recapitation” method allows msprime to be used to extremely efficiently add a “neutral burn-in” history to a completed SLiM simulation of a selective sweep, by coalescing the simulation’s initial population backward in time.

## Examples

Examples were executed on a MacBook Pro (2.9 GHz Intel Core i7, 16 GB RAM) running macOS 10.13.5, using Python 3.4.8, R 3.5.0, SLiM 3.1, msprime 0.6.1, and pyslim 0.1. Reported times were measured with the Python `timeit` package. Peak memory usage for SLiM runs was assessed with SLiM’s `-m` command-line option. The timing comparison (Figure 2) was executed on the same hardware, with macOS 10.13.4, R 3.4.3, SLiM 3.0, and msprime 0.6.0, using the Un*x tool `/usr/bin/time` for timing (summing the reported user time and system time); we believe the times measured would not change significantly with the newer software versions. The full source code for the examples and timing tests, including timing and plotting code that is omitted here, may be found at https://github.com/bhaller/SLiMTreeSeqPub. These examples use the `matplotlib` (Hunter, 2007) and `numpy` (Oliphant, 2006) packages for Python.

**Figure 2.**
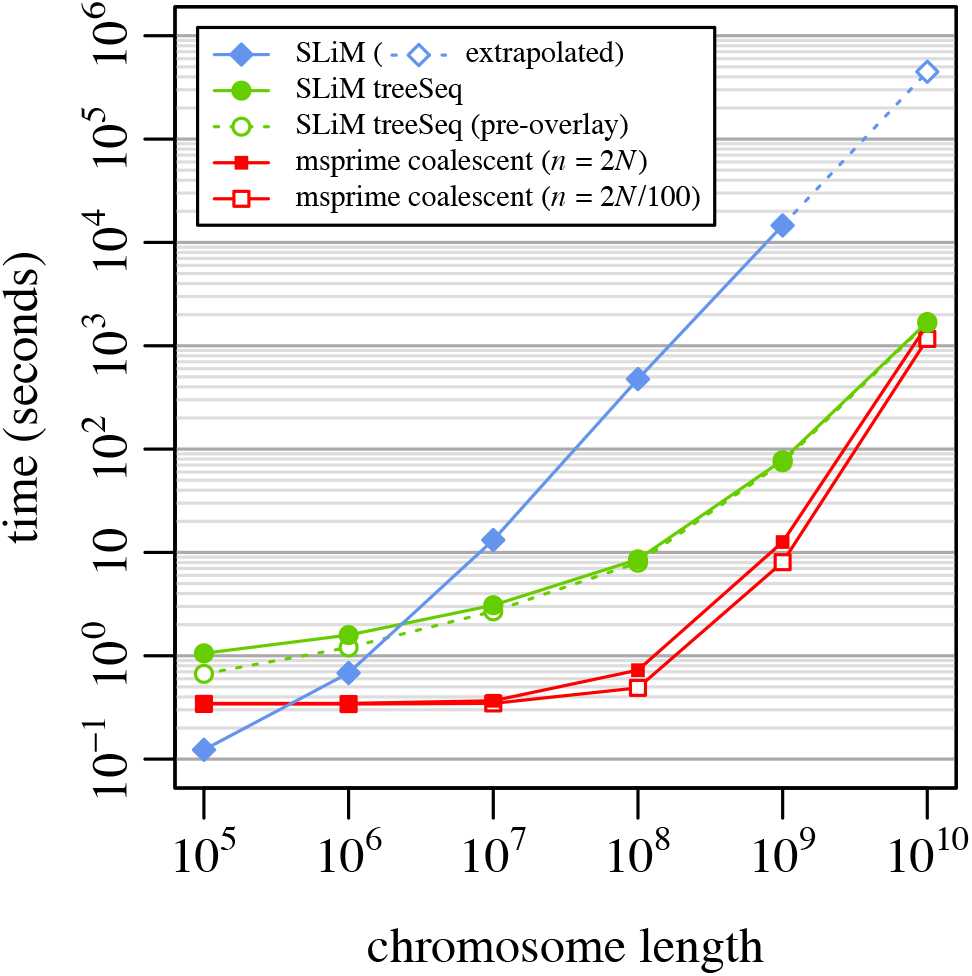
A speed comparison between SLiM without tree-sequence recording, SLiM with tree-sequence recording and mutation overlay, and msprime’s coalescent simulation for a simple neutral model (Example 1; see text for model description). Each point represents the mean runtime across 10 replicates using different random number seeds; bars showing standard error of the mean would be smaller than the size of the plotted points in all cases. Runs for SLiM without tree-sequence recording (filled blue diamonds) were not conducted for *L* = 10^10^ because the memory usage was prohibitive, so a linear extrapolation is shown (hollow blue diamond). Runs for SLiM with tree-sequence recording and mutation overlay (filled green circles) are subdivided here to show the runtime for SLiM alone, prior to mutation overlay (hollow green circles), illustrating that the time for mutation overlay is negligible. The runtimes for the msprime coalescent for a full population sample of *n* = 2*N* = 1000 (filled red squares) and for a sample of size *n* = 2*N*/100 = 10 (hollow red squares) are both shown. Note that the *x* and *y* axes are both on a log scale.

### Example I: A simple neutral model

Our first example is a model of a neutrally evolving chromosome of length *L* = 10^8^ base positions, with uniform mutation rate *μ* = 10^−7^ and recombination rate *r* = 10^−8^ (both expressed as the event probability per base per generation), in a panmictic diploid population of size *N* = 500, running for a duration of 10*N* = 5000 non-overlapping generations. The SLiM configuration script for this basic model is very simple:

~~~
         initialize() {
              initializeMutationRate(1e-7);
              initializeMutationType("m1", 0.5, "f", 0.0);
              initializeGenomicElementType("g1", m1, 1.0);
              initializeGenomicElement(g1, 0, 1e8-1); initializeRecombinationRate(1e-8);
         }
         1 {
              sim.addSubpop("p1", 500);
         }
         5000 late() {
              sim.outputFull("ex1_noTS.slimbinary", binary=T);
         }
~~~

This sets up a single “genomic element” spanning the full length of the chromosome, with neutral mutations of type m1 generated at the desired rate, and with the desired recombination rate. In generation 1 a new subpopulation of the desired size is created, and the model runs to generation 5000, after which it outputs the full simulation state. The SLiM manual provides additional explanation of these concepts (Haller and Messer, 2016). This model took 211.9 seconds to run, and reached a peak memory usage of 443.8 MB.

Tree-sequence recording can easily be enabled for this model with a call to

~~~
         initializeTreeSeq():
              initialize() {
              initializeTreeSeq();
              initializeMutationRate(0);
              initializeMutationType("m1", 0.5, "f", 0.0);
              initializeGenomicElementType("g1", m1, 1.0);
              initializeGenomicElement(g1, 0, 1e8-1); initializeRecombinationRate(1e-8);
         }
         1 {
              sim.addSubpop("p1", 500);
         5000 late() {
              sim.treeSeqOutput("ex1_TS.trees");
         }
~~~

Note that we have now also set the mutation rate to zero; SLiM no longer needs to model neutral mutations because they can be overlaid in a later step more efficiently. A `.trees` file is output at the end of the run, instead of calling SLiM’s `outputFull()` method, so that the recorded tree sequence is preserved. In all other respects these models are identical. This is typical of adapting a SLiM model to use tree-sequence recording: in general, the aim is to remove the modeling of neutral mutations while preserving other aspects of the model verbatim.

After simulation has completed, neutral mutations are overlaid upon the saved tree sequence. The full model – running the SLiM model and then doing the final mutation overlay step – can be executed with a simple Python script:

~~~
         import subprocess, msprime, pyslim
         # Run the SLiM model
         subprocess.check_output(["slim", "-m", "-s", "0", "ex1_TS.slim"])
         # Overlay neutral mutations
         ts = pyslim.load("ex1_TS.trees")
         mutated = msprime.mutate(ts, rate=1e-7, random_seed=1, keep=True)
         mutated.dump("ex1_TS_overlaid.trees")
~~~

This script uses the msprime Python package to overlay neutral mutations upon the recorded tree sequence. The result is precisely the same, statistically, as if the neutral mutations were included in the forward simulation, except that the vast majority of the bookkeeping work in each generation is avoided because mutations only need to be overlaid upon the ancestral genomic regions that persisted to the end of the simulation.

Note that pyslim is used to load the `.trees` file; this package provides a bridge between SLiM and msprime, and should generally be used to load and save `.trees` files in Python if the files are coming from or going to SLiM. The pyslim package extends the msprime tree sequence class by adding support for SLiM’s metadata annotations to the tree sequence, providing an interface for reading or modifying that metadata as well as for generating SLiM-compliant `.trees` files that contain the required metadata. The `.trees` files output by SliM can be read directly by msprime, but the returned object will have reduced functionality compared to those returned by pyslim.

The total time to execute this Python code is 4.37 seconds, almost 50 times faster than the model without tree-sequence recording. Most of the runtime (4.09 seconds) is spent running the SLiM model; the final mutation overlay by msprime is extremely fast. The peak memory usage during the SLiM run is 145.8 MB, less than one-third of the memory usage of the model without tree-sequence recording. Tree-sequence recording can often reduce memory usage, since the tree sequence data structure is quite compact compared to SLiM’s in-memory representation of the neutral mutations that would be segregating in such a model. Tree sequences are also very compact on disk; the final `.trees` file here, with mutations overlaid, takes about 8.9 MB, as compared to 84.2 MB for the `ex1_noTS.slimbinary` file from the SLiM model without tree-sequence recording, 559 MB for a Newick file, and 366 MB for a VCF file – even though the `.trees` file contains ancestry information not included by the SLiM and VCF formats. A VCF file containing the sequences of the final generation can be produced from a `.trees` file with msprime’s `write_vcf()` method, but the ancestry information is lost.

The speedup produced by this tree-sequence recording method can vary dramatically depending upon the details of the simulation; all of the work to track neutral mutations is eliminated, but new work is added involving the recording of all the recombination events that go into producing the tree sequence. In general, the largest speedup will be observed with very long chromosomes with many neutral mutations when the recombination rate is not too high; indeed, when modeling a very short chromosome the overhead of tree-sequence recording can outweigh the savings from omitting neutral mutations (see Discussion).

To further illustrate the performance benefits of tree-sequence recording, we conducted a set of timing comparisons between SLiM without tree-sequence recording, SLiM with tree-sequence recording, and msprime’s coalescent simulation method. These comparisons involved essentially the same model as shown above: a neutral panmictic model of diploids with non-overlapping generations, with a population size *N* = 500, recombination rate *r* = 10^−8^ per base position per generation, and mutation rate *μ* = 10^−7^ per base position per generation. The chromosome length *L* was varied over {10^5^, 10^6^, 10^7^, 10^8^, 10^9^, 10^10^}, with ten runs of each model at each value of *L* using different random seeds. The number of generations varied with *L* (details below). The msprime coalescent was run both with a final haploid sample size *n* equal to the full population size (*n* = 2*N*), and with a much smaller sample size (*n* = 2*N*/100); in both cases, *N*_e_ = *N* was used. To verify that tree-sequence recording produced results equivalent to the coalescent, we checked that the mean TMRCAs for the *L* = 10^10^ runs for the two methods did not differ significantly (*p* = 0.7791).

The average runtimes obtained are shown in Figure 2. As *L* increased, the benefit of tree-sequence recording compared to SLiM without tree-sequence recording became increasingly large, topping out at a performance improvement of more than two orders of magnitude for *L* = 10^9^ and *L* = 10^10^. Coalescent simulations with msprime were much faster than the tree-sequence recording method, as expected, except at *L* = 10^10^, where msprime’s speed was comparable to that of SLiM with tree-sequence recording. It appears that SLiM with tree-sequence recording would be faster for *L* larger than 10^10^. The number of events the coalescent must simulate is quadratic in *L*, empirically, but with a small leading coefficient such that msprime is quite fast even for reasonably large chromosome sizes (Kelleher et al. 2016). With very large values of *L*, however, this O(*L*^2^) term begins to dominate and SLiM with tree-sequence recording becomes faster. This may be chiefly of theoretical interest, since *L* = 10^10^ is already a very long chromosome (approximately three times the length of the full human genome). It is also noteworthy that the msprime coalescent is only marginally faster for a sample of *n* = 2*N*/100 than for a full population sample of *n* = 2*N*; as more samples are added to a gene tree, the new samples tend to attach to already existing branches quite quickly (Kingman, 1982).

Although the coalescent remains an order of magnitude faster for most practical purposes, it can only be used in a few simple scenarios such as this; for models that require forward simulation, tree-sequence recording offers large performance benefits over more traditional forward simulation techniques. It is also worth noting that the coalescent is only an approximation of the Wright–Fisher model, and will diverge from it under certain conditions (Wakeley et al., 2012; Bhaskar et al., 2014) – one such condition being a sample size that is no longer small compared to the population size, as is the case for our *n* = 2*N* msprime runs here. Forward simulation may therefore be preferable in order to obtain exact results under such conditions.

**How long do we run it?** In general, it is desirable to run forward-time simulations “until convergence” – until the effects of the starting configuration are forgotten. This occurs (in most situations) when all genealogical trees have coalesced, meaning that at every position in the genome a common ancestor to the entire final generation has appeared. In practice, models are often run for 10*N* generations, a rule of thumb that is thought to suffice in most cases. However, this is a thorny problem: longer chromosomes tend to require longer for coalescence, simply because with more sites it is more likely that coalescence takes exceptionally long at some site. In the simulations of Figure 2, we ran each simulation for the expected number of generations required for coalescence at that value of *L*, which increased linearly with log(*L*), from about 3*N* for *L* = 1e5 to 15*N* for *L* = 1e10. This sufficed to make the comparison between SLiM and msprime “fair”, but a better practical solution, recapitation, will be shown in Example 4. We determined the expected number of generations empirically by running the same model 500 times at each value of *L* with “coalescence detection” enabled (by passing checkCoalescence=T to initializeTreeSeq()). The mean and other summary statistics for each value of *L* (Table S1) agree with expectations from extreme value theory (Berman, 1964), with the expected time until coalescence growing roughly as 1000 log(*L*) − 10000.

### Example II: Background selection

Our second example is a model of background selection, a term which describes the effect that purifying selection against deleterious mutations imposes on genetic variation at linked sites. Such purifying selection should be particularly common in genic regions, where many genomic positions should be subject to selective constraints. This background selection, like many types of linked selection more generally, is expected to produce a “dip in diversity” in the surrounding non-coding regions, with a signature of decreasing genetic diversity with decreasing distance to the nearest gene (Charlesworth et al. 1993; Hudson 1994; Sattath et al., 2011; Elyashiv et al., 2016). Here is a SLiM model that uses tree-sequence recording to model this scenario:

~~~
         initialize() {
             defineConstant("N", 10000); // pop size
             defineConstant("L", 1e8); // total chromosome length defineConstant("L0", 200e3); // between genes
             defineConstant("L1", 1e3); // gene length
             initializeTreeSeq();
             initializeMutationRate(1e-7); initializeRecombinationRate(1e-8, L-1);
             initializeMutationType("m2", 0.5, "g", -(5/N), 1.0);
             initializeGenomicElementType("g2", m2, 1.0);
             for (start in seq(from=L0, to=L-(L0+L1), by=(L0+L1)))
             initializeGenomicElement(g2, start, (start+L1)-1);
         }
         1 {
             sim.addSubpop("p1", N);
             sim.rescheduleScriptBlock(s1, 10*N, 10*N);
         }
         s1 10 late() {
             sim.treeSeqOutput("ex2_TS.trees");
         }
~~~

This model sets up a chromosome that consists of genes of length *L*_1_ = 1 kb, separated by non-coding regions of length *L*_0_ = 200 kb. The total chromosome length is *L* = 10^8^ bases, and 496 genes fit within it. The model uses a mutation rate of *μ* = 10^−7^ for deleterious mutations that can arise within the genes; no other mutations are modeled. The deleterious mutations are given selection coefficients drawn from a Gamma distribution with mean −5/*N* and shape parameter *α* = 1 (modeling a scenario of moderately deleterious mutations with 2*Ns* = −10 on average). We assume co-dominance with *h* = 0.5. A population of size *N* = 10000 is started in generation 1, and the model runs until generation *G* = 10*N* (the output event, s1, is rescheduled dynamically to that generation).

We can run this model and then conduct post-run analysis with a Python script:

~~~
         import os, subprocess, msprime, statistics, pyslim
         import matplotlib.pyplot as plt
         import numpy as np
         # Run the SLiM model and load the resulting .trees file
         subprocess.check_output(["slim", "-m", "-s", "0", "ex2_TS.slim"])
         ts = pyslim.load("ex2_TS.trees").simplify()
         # Measure the tree height at each base position
         height_for_pos = np.zeros(int(ts.sequence_length))
         for tree in ts.trees():
              mean_height = statistics.mean([tree.time(root) for root in tree.roots])
              left, right = map(int, tree.interval)
              height_for_pos[left: right] = mean_height
         # Convert heights along the chromosome into heights at distances from a gene
         height_for_pos = height_for_pos - np.min(height_for_pos)
         L, L0, L1 = int(1e8), int(200e3), int(1e3)
         gene_starts = np.arange(L0, L - (L0 + L1) + 1, L0 + L1)
         gene_ends = gene_starts + L1 - 1
         max_distance = L0 // 4
              height_for_left_distance = np.zeros(max_distance)
              height_for_right_distance = np.zeros(max_distance)
         for d in range(max_distance):
                  height_for_left_distance[d] = np.mean(height_for_pos[gene_starts - d - 1])
              height_for_right_distance[d] = np.mean(height_for_pos[gene_ends + d + 1])
              height_for_distance = np.hstack([height_for_left_distance[::-1],
              height_for_right_distance])
         distances = np.hstack([np.arange(-max_distance, 0), np.arange(1, max_distance + 1)])
         # Make a simple plot plt.plot(distances, height_for_distance)
         plt.show()
~~~

The first line after the import statement runs the SLiM model; this took 15643 seconds (4.35 hours) to execute. This is not short – it is still a fairly complex model! – but it is far shorter than the alternative, a SLiM model without tree-sequence recording and including neutral mutations in the non-coding regions. That alternative model would take ~83 hours, by extrapolation – probably a conservative estimate, since the model had not yet reached mutation–selection balance and was still slowing down when its timing was measured. The use of tree-sequence recording here results, then, in a relatively modest speedup of 19 times. This makes sense, since the model with tree-sequence recording still must keep track of a very large number of segregating deleterious mutations. However, it is worth noting that the final result from this alternative model would provide far less statistical power, since inference from it would be based only upon the observed pattern of neutral mutations in one run, rather than the actual pattern of ancestry at each chromosome position; to provide the same power, this alternative model would likely have to be run many times or use a much higher mutation rate. If more performance gains were needed, the model could perhaps be rescaled as well (see Discussion).

The rest of the code conducts post-run analyses. First, the `.trees` file from the SLiM run is read in with `pyslim.load()` as in the previous example; here, however, we call `simplify()` (Kelleher et al. 2018) upon the loaded tree sequence, which requires some explanation. SLiM automatically retains, in the tree sequence, nodes corresponding to the original ancestors of each subpopulation that was created with `addSubpop()`. This is done for various reasons, including allowing ancestry to be more easily traced and enabling recapitation (see Example 4). When SLiM saves a `.trees` file, these ancestors are present in the tree sequence but are not marked as “samples”, and will therefore disappear after a `simplify()` operation. In many cases these ancestors are harmless, as in Example 1; in fact, in Example 1, calling `simplify()` to remove them would mean that mutations would be overlaid only back to the point of coalescence, rather than to the beginning of forward simulation. Here, however, since we want to measure the heights of trees in the tree sequence, these ancestors would complicate things for us; all trees would be rooted in those ancestors, at the beginning of forward simulation. We therefore call `simplify()` to remove them (when the model has coalesced below them; they are retained when still in use by the tree sequence). Example 4 will delve into this matter further.

Next, a vector containing the mean tree height at each base position (`height_for_pos`) is constructed by walking through the tree sequence to find the set of trees representing the ancestry of every individual in the final generation at a given position. The mean tree height is a metric of the time to the most recent common ancestor at a given base position, and thus of diversity at that base position; background selection will tend to reduce the mean tree height, thereby lowering the expected levels of diversity at a locus.

An aside: there can be a set of trees for a given position, rather than just a single tree, if the forward simulation was not run sufficiently long for coalescence to have occurred at every position in the genome. In msprime this is modelled by allowing trees to have multiple roots. Each root represents the most recent common ancestor of some subset of the extant population at that location in the genome; if coalescence has not occurred, then the final population should still contain genetic variation that was segregating in the initial population, since different individuals inherit from different roots of the ancestry tree. Since the model here ran for 10*N* generations, we can hope that it has coalesced at most or all positions; but unless a model is explicitly run out to coalescence (or recapitated), it is always possible that multiple roots will exist, and so robust code ought to handle that case by looping over the roots for each tree as we do here.

These mean tree heights along the chromosome are then converted to mean tree heights at distances from the nearest gene (`height_for_distance`), taking into account the somewhat complex genetic structure of the model. Finally, the relationship between distance to the nearest gene and tree height is plotted. These analyses took 12.39 seconds to complete. Note that neutral mutations were never simulated at all; the analysis is based upon the tree sequence itself, not upon the distribution of neutral mutations.

A plot of the results can be seen in Figure 3, showing the well-known “dip in diversity” realized here through simulation. As the distance to the nearest gene decreases, diversity dips due to the background selection exerted by selection against deleterious mutations within the gene.

**Figure 3.**
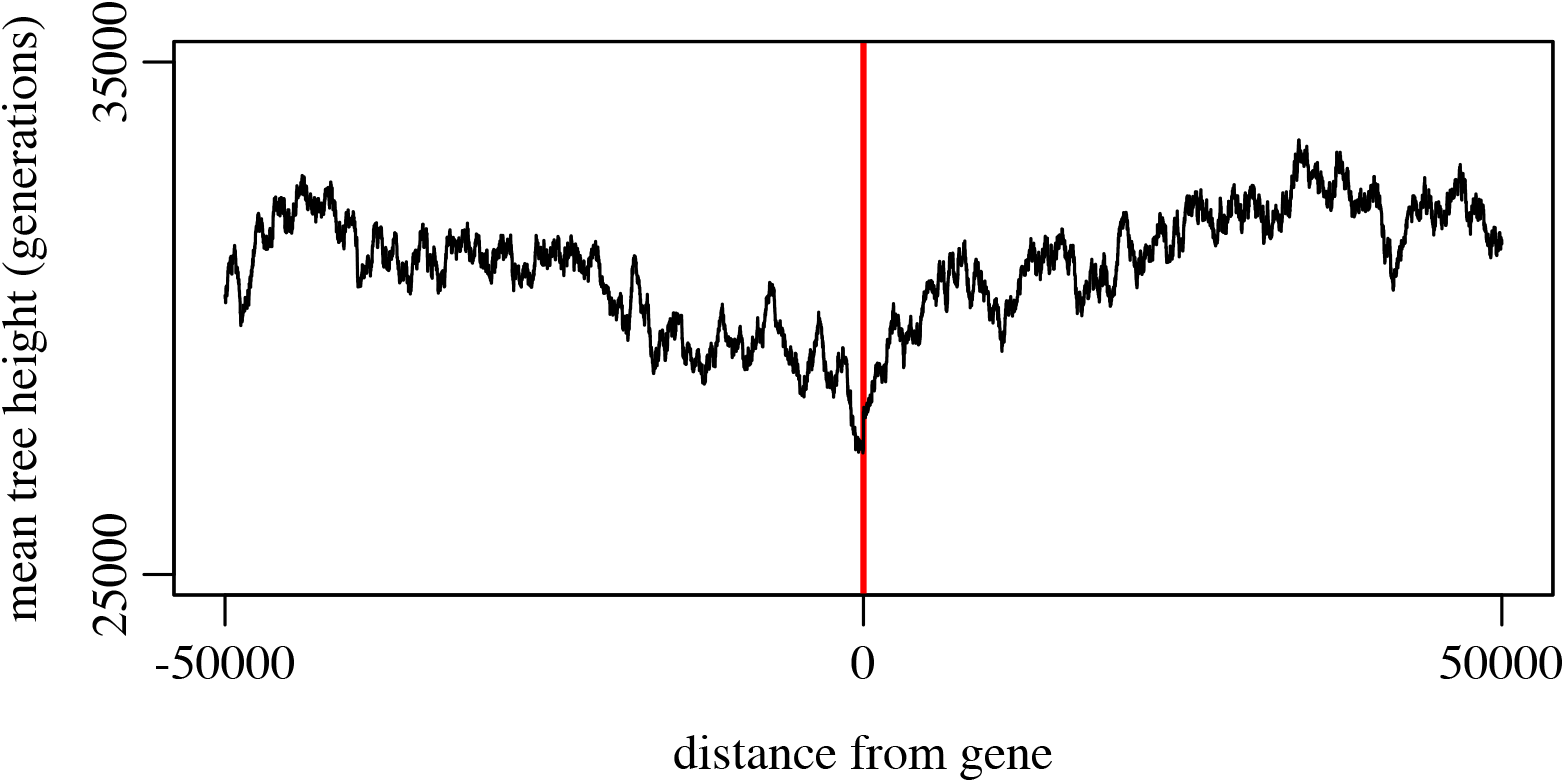
Mean diversity (as measured by mean tree height) as a function of distance from the nearest gene (Example 2). The center of the x-axis (red line) represents a distance of zero, immediately adjacent to a gene; moving away from the x-axis center to the left/right represents moving away from the nearest gene to the left/right respectively. The pattern of decreased diversity near a gene is the “dip in diversity” due to background selection.

### Example III: True local ancestry mapping

Our third example involves mapping the true local ancestry at every position along a chromosome in a two-subpopulation admixture model with adaptive introgression at two partially linked loci. This is an important dynamic in all sorts of biological systems, from human–Neanderthal admixture to hybrid zones between divergent bird populations; one often wishes to be able to find which ancestral population each chromosomal region traces back to. The SLiM model looks like this:

~~~
         initialize() { defineConstant("L", 1e8);
                initializeTreeSeq(); initializeMutationRate(0);
                initializeMutationType("m1", 0.5, "f", 0.1);
                initializeGenomicElementType("g1", m1, 1.0);
                initializeGenomicElement(g1, 0, L-1);
                initializeRecombinationRate(1e-8);
         }
         1 late() { sim.addSubpop("p1", 500);
                sim.addSubpop("p2", 500);
                sim.treeSeqRememberIndividuals(sim.subpopulations.individuals);
                p1.genomes.addNewDrawnMutation(m1, asInteger(L * 0.2));
                p2.genomes.addNewDrawnMutation(m1, asInteger(L * 0.8));
                sim.addSubpop("p3", 1000); p3.setMigrationRates(c(p1, p2), c(0.5, 0.5));
         }
                2 late() {
                p3.setMigrationRates(c(p1, p2), c(0.0, 0.0));
                p1.setSubpopulationSize(0);
                p2.setSubpopulationSize(0);
         }
         2: late() {
                if (sim.mutationsOfType(m1).size() == 0)
                {
                sim.treeSeqOutput("ex3_TS.trees");
                sim.simulationFinished();
         }
         }
                10000 late() {
                stop("Did not reach fixation of beneficial alleles.");
         }
~~~

The `initialize()` callback sets up tree-sequence recording with a mutation rate of *μ* = 0 and a recombination rate of *r* = 10^−8^ along a chromosome of length *L* = 10^8^. Although the mutation rate is zero, a mutation type m1 is defined representing beneficial mutations with a selection coefficient of *s* = 0.1; mutations of this type will be added in generation 1.

In generation 1 we create two subpopulations, p1 and p2, of 500 individuals each; these are the original subpopulations that will admix. We tell SLiM to remember these individuals forever as ancestors in the tree sequence, with treeSeqRememberIndividuals(), because we want them to act as the roots of all recorded trees so that we can establish local ancestry using them. Note that this is not strictly necessary, since (as discussed in Example 2) SLiM automatically retains the root ancestors for each population; we could rely upon that, and we would be fine as long as we did not `simplify()` after loading the tree sequence in Python. The use of treeSeqRememberIndividuals() has been shown here for purposes of illustration, however, since some models may wish to remember non-root individuals for analysis. Next, we add a beneficial mutation at 0.2*L* in p1, and another at 0.8*L* in p2; the expectation is that by the end of the run all individuals will be recombinants that carry both of these mutations. Finally, we create subpopulation p3 and tell SLiM that it will be composed entirely of migrants from p1 and p2 in equal measure.

By the end of generation 2, subpopulation p3 has received its offspring generation from p1 and p2 as intended, so we can now remove p1 and p2 from the model and allow p3 to evolve. At this stage, all individuals in p3 are still unmixed, having been generated from parents in either p1 or p2, but beginning in generation 3 they will start to mix.

Finally, we have some output and termination code. If both m1 mutations fix, they are converted to `Substitution` objects by SLiM, and when that is detected the model writes out a final `.trees` file and terminates. If we reach generation 10000 without that happening, the admixture failed, and we stop with an error. This model is conceptually similar to recipe 13.9 in the SLiM manual (Haller and Messer, 2016), but has been converted to use tree-sequence recording, so you can refer to the manual’s recipe for additional commentary.

We can run this model from a Python script and do post-run analysis, as we did in Example 2:

~~~
         import os, subprocess, msprime, pyslim
         import matplotlib.pyplot as plt
         import numpy as np
         # Run the SLiM model and load the resulting .trees file
         subprocess.check_output(["slim", "-m", "-s", "0", "ex3_TS.slim"])
         ts = pyslim.load("ex3_TS.trees").simplify()
         # Assess the true local ancestry at each base position
         breaks = np.zeros(ts.num_trees + 1)
         ancestry = np.zeros(ts.num_trees + 1)
         for tree in ts.trees(sample_counts=True):
              subpop_sum, subpop_weights = 0, 0
              for root in tree.roots:
                   leaves_count = tree.num_samples(root) - 1 # subtract one for the root
                   subpop_sum += tree.population(root) * leaves_count
                   subpop_weights += leaves_count breaks[tree.index] = tree.interval[0]
                   ancestry[tree.index] = subpop_sum / subpop_weights
              breaks[-1] = ts.sequence_length
              ancestry[-1] = ancestry[-2]
              # Make a simple plot
         plt.plot(breaks, ancestry)
         plt.show()
~~~

The first line after the import statements runs the SLiM model, which completes in just 0.416 seconds, with peak memory usage of 55.6 MB; since it tracks only two mutations, and typically terminates by generation 150 or so, it is very quick.

The equivalent SLiM model to achieve true local ancestry mapping without tree-sequence recording has to model a mutation at each base position, as can be seen in recipe 13.9 in the SLiM manual (Haller and Messer, 2016). A direct comparison is not possible, because recipe 13.9 scaled up to a chromosome length of *L* = 10^8^ would take an estimated 7.2 days to run, and worse, would require 8.1 TB (terabytes) of memory. Those estimates are derived from the pattern of performance observed for recipe 13.9 with *L* = 5×10^5^, *L* = 10^6^, and *L* = 2×10^6^ (the upper limit on our test machine due to memory usage), extrapolated out to *L* = 10^8^. Implementing this model with tree-sequence recording therefore reduces the runtime by a factor of more than 1.35 million, and reduces the memory usage by a factor of more than 160,000.

Similar to Example 2, the post-run analysis walks through the tree sequence, but in this case, computes the mean true local ancestry (the fractional ancestry from subpopulation p1 versus p2) for each tree. This is done by finding the roots for the tree, assessing the subpopulations of origin of those root individuals, and averaging those together weighted by the number of descendants from each root. A simple plot is then produced. In this example, the analysis took 62.2 seconds; the analysis runtime is relatively long because the trees here typically have many roots, so the inner loop is executed a great many times.

The final plot of true local ancestry by chromosome position is shown in Figure 4. The mean true local ancestry at the points where the beneficial mutations were introduced into p1 and p2 has to be 100% p1 and 100% p2, respectively, since both beneficial mutations fixed by the end of the run. At other points along the genome there is more variation, but with a general pattern of being more completely admixed at the chromosome ends and middle, with gradations toward the absolute p1 and p2 points. Since this is a single run of the model, the pattern is quite stochastic; an average across many runs of this model could produce a smooth plot if desired, and since it takes only a couple of minutes to execute the pipeline here, that would be very quick to do. This method of calculating true local ancestry could be used by any SLiM model with tree-sequence recording, so models with more complex demography, under any scenario of selection and mating, with any recombination map, etc., could just as easily be explored.

**Figure 4.**
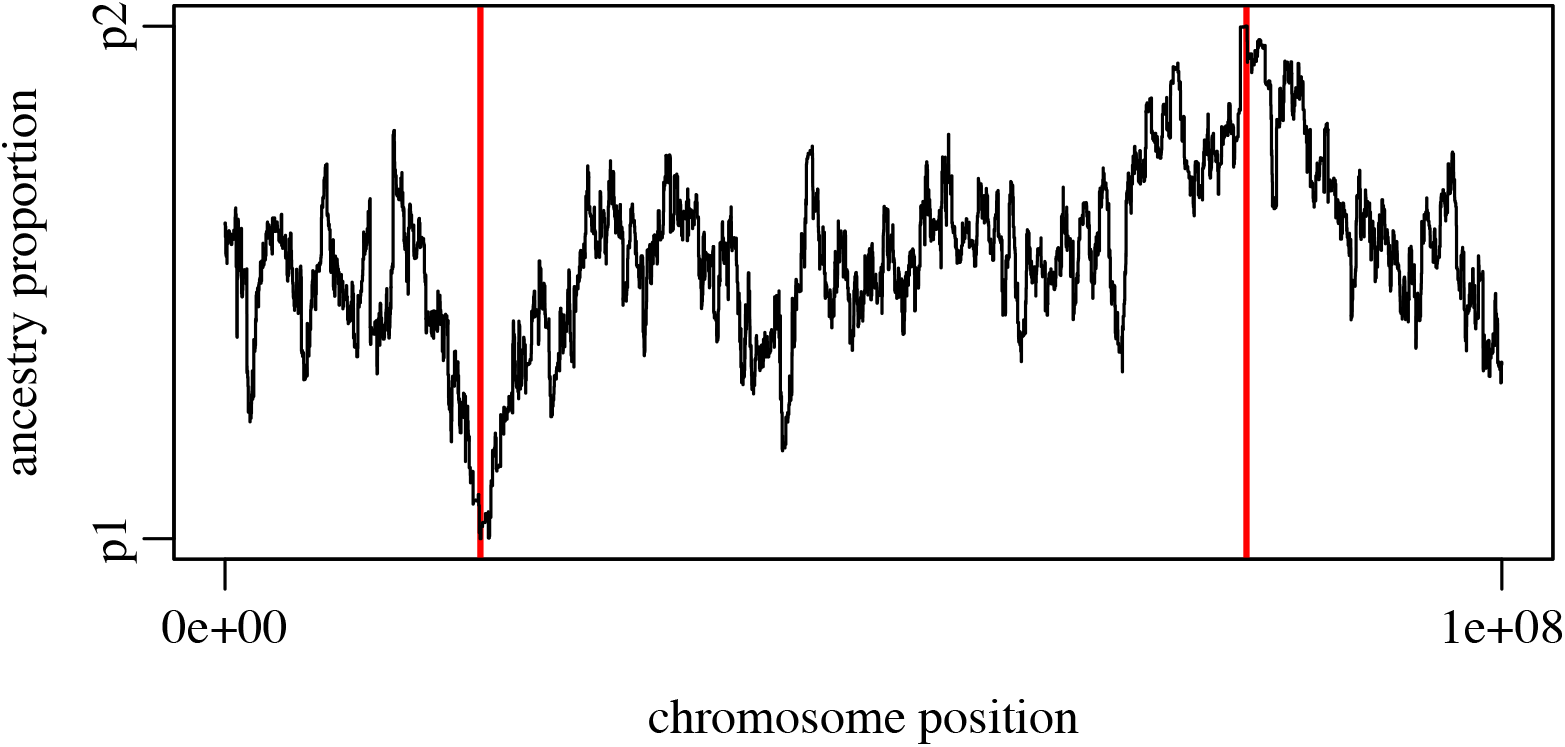
Mean true local ancestry at each position along the chromosome (Example 3). The red vertical bars indicate the positions at which beneficial mutations were originally introduced into p1 and p2. The beneficial mutations, which both fixed, are points where the true local ancestry is 100% p1 or p2. True local ancestry regresses toward equal admixture with increasing distance from those fixed points.

### Example IV: Neutral burn-in for a non-neutral model

Our final example illustrates a solution to the problem of neutral burn-in. In many applications one wishes to execute a non-neutral forward simulation beginning with an equilibrium amount of extant neutral genetic diversity, and the simulation needed to generate that pre-existing diversity, typically called the model “burn-in”, can take quite a long time – often much longer than it takes to execute the non-neutral portion of the simulation. For a model with a long chromosome or large population size, this burn-in can be so long as to limit the practical scale of the simulations that can be conducted. One solution to this is a “hybrid” approach, in which a forward simulation is initialized with the result of a (much faster) coalescent simulation (similar to Bhaskar 2014). This is now possible using tree sequences in SLiM, but we go a step further: even a great deal of the work done in a coalescent simulation of this burn-in period is unnecessary. All of the genealogical branches that go extinct are irrelevant; all that matters are those segments of ancestral genomes from which the final generation inherits. With tree-sequence recording, one can simulate only the histories of those segments, saving an immense amount of computation relative to a forward-time burn-in simulation.

Here we will look at a fairly large model (*N* = 10^5^; *L* = 10^6^) that evolves under neutral dynamics until coalescence (the neutral burn-in), after which follows some relatively brief non-neutral dynamics (a selective sweep). Running the burn-in period for this model in SLiM would take an exceedingly long time, given the scale of the model, as we will see below. A better idea is to use what we call “recapitation”: we can run the SLiM model forward from an initial state that conceptually follows burn-in, and then use msprime to generate *after the fact* the coalescent history for the initial individuals of the forward simulation. This can be done without simulating neutral mutations, but if neutral mutations are desired as an end product of the simulation, they can be overlaid at the end as in Example 1.

We begin with the SLiM model, which simulates the introduction and sweep to fixation of a beneficial mutation. For simplicity, we will select a run of the model that happens to result in fixation, rather than using a recipe that is conditional upon fixation; the random number seed specified in the Python script below should produce that outcome. The SLiM model:

~~~
         initialize() { initializeTreeSeq(simplificationRatio=INF);
              initializeMutationRate(0); initializeMutationType("m2", 0.5, "f", 1.0);
              m2.convertToSubstitution = F;
              initializeGenomicElementType("g1", m2, 1);
              initializeGenomicElement(g1, 0, 1e6 - 1);
              initializeRecombinationRate(3e-10);
         }
         1 late() {
              sim.addSubpop("p1", 100000);
         }
              100 late() {
              sample(p1.genomes, 1).addNewDrawnMutation(m2, 5e5);
         }
         100:10000 late() {
              mut = sim.mutationsOfType(m2);
              if (mut.size() != 1)
              stop(sim.generation + ": LOST");
              else if (sum(sim.mutationFrequencies(NULL, mut)) == 1.0)
         {
              sim.treeSeqOutput("ex4_TS_decap.trees");
              sim.simulationFinished();
         }
         }
~~~

This specifies a simple model with population size *N* = 10^5^ diploid individuals, chromosome length *L* = 10^6^ base positions, and a recombination rate of *r* = 3×10^−10^ per base position per generation, without mutation. It runs to generation 100 and then introduces the sweep mutation (the delay before introduction is just to provide separation between the simulation start and the start of the sweep in the plot produced below). When the sweep mutation is found to have fixed, it then outputs a `.trees` file and stops. It specifies an infinite “simplification ratio” in the call to `initializeTreeSeq()` so that simplification happens only once, at the point when the `.trees` file is written out at the end; with this large of a model simplification takes a significant amount of time, so this optional setting speeds the model up somewhat at the price of a higher peak memory footprint.

As in previous examples, we will run this from a Python script that does post-run analysis:

~~~
         import os, subprocess, msprime, pyslim
         import numpy as np
         import matplotlib.pyplot as plt
         # Run the SLiM model and load the resulting .trees file subprocess.check_output(["slim", "-m", "-s", "2", "ex4_TS.slim"])
         ts = pyslim.load("ex4_TS_decap.trees") # no simplify!
         # Calculate tree heights
         def tree_heights(ts):
             heights = np.zeros(ts.num_trees + 1)
             for tree in ts.trees():
                  if tree.num_roots > 1: # not fully coalesced heights[tree.index] = ts.slim_generation
                  else:
                  root_children = tree.children(tree.root)
                  real_root = tree.root if len(root_children) > 1 else root_children[0]
                  heights[tree.index] = tree.time(real_root)
             heights[-1] = heights[-2] # repeat the last entry for plotting with step
            return heights
         # Plot tree heights before recapitation
         breakpoints = list(ts.breakpoints())
         heights = tree_heights(ts)
         plt.step(breakpoints, heights, where=’post’)
         plt.show()
         # Recapitate
         >recap = ts.recapitate(recombination_rate=3e-10, Ne=1e5, random_seed=1)
         recap.dump("ex4_TS_recap.trees")
         # Plot tree heights after recapitation
         breakpoints = list(recap.breakpoints())
         heights = tree_heights(recap) plt.step(breakpoints, heights, where=’post’)
         plt.show()
~~~

After the import, we run the SLiM model (which takes 46.05 seconds) and load the `.trees` file it saves out. We then immediately make a plot of mean tree heights along the chromosome. This is similar to what we did in Example 2, but here it requires some extra finesse because we did not simplify the tree sequence after loading it as we did then. To perform recapitation, we cannot first simplify – we need the ancestral individuals that started the SLiM simulation to remain in the tree sequence, so that recapitation can build upon them correctly. For this reason, every root in the loaded tree sequence has the same time, corresponding to the beginning of the forward simulation. The code in the `tree_heights()` function corrects for that, getting the height of the child of the root if the forward simulation has coalesced below the original ancestor. This provides the red line in Figure 5, showing that the area immediately around the introduced mutation has coalesced at the time of the introduction (due to hitchhiking), but that the remainder of the chromosome has not yet coalesced and thus has a tree height corresponding to the start of forward simulation. These uncoalesced plateaus are what we will fix with recapitation.

**Figure 5.**
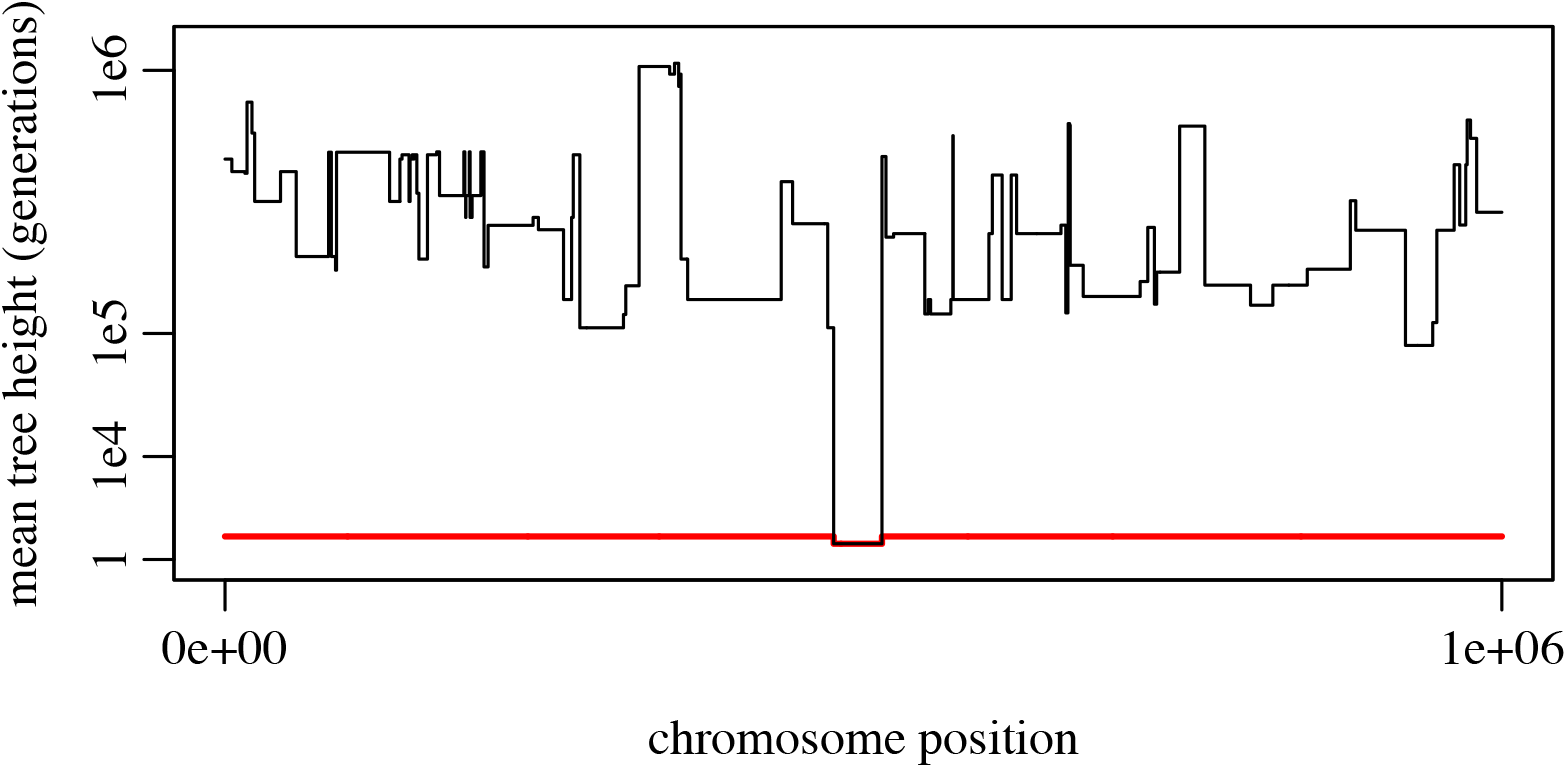
Mean tree height (on a cube-root-scaled *y*-axis) at each position along the chromosome, before and after recapitation (Example 4). The red line shows mean tree heights prior to recapitation; the region surrounding the introduced sweep mutation coalesces at the start of the sweep, whereas the plateaus outside that region are uncoalesced and have a height corresponding to the start of forward simulation (100 generations earlier). The black line shows heights after recapitation; the uncoalesced plateaus have now been coalesced backward in time, producing tree heights as long as a million generations in the past.

The next step, then, is to perform the recapitation. This process works backwards from the tree sequence information recorded by SLiM, constructing a full coalescent history for all of the individuals alive at the end of the run. Since the non-neutral dynamics eliminated much of the genetic diversity from the population as it existed at the beginning of forward simulation, this coalescence requires very little work – much less than even a normal coalescent simulation for this population size would require. In the example run discussed here, the process took 0.41 seconds. If neutral mutations are desired, they can then be overlaid on the recapitated tree sequence following the method of Example 1; that code is not shown again here, but that operation took another 0.58 seconds (with *μ* = 10^−7^).

Finally, we plot the mean tree heights for the recapitated tree sequence; this produces the black line in Figure 5. The uncoalesced plateaus have now coalesced to times as far as a million generations in the past. This plot nicely illustrates the classical sweep pattern in which regions closer to the position of the sweep tend to coalesce more recently, due to hitchhiking, than regions farther away (Maynard-Smith and Haigh, 1974).

Simulating the neutral burn-in period in SLiM instead, with neutral mutations occurring at a rate of *μ* = 10^−7^, would take an estimated 114.7 hours (from extrapolation; this is a very conservative estimate since the model was nowhere near mutation–drift balance when times were measured). Recapitation and neutral mutation overlay, with a total time of 0.99 seconds, therefore sped up the burn-in process in this example by more than 400,000 times.

Recapitation is clearly much faster than conducting burn-in with forward simulation, then; it should be faster than a rescaled forward simulation model too (since rescaling can generally not be taken that far without introducing problematic artifacts; see Discussion), and faster even than constructing the burn-in state with the coalescent (since recapitation is based upon the coalescent but handles far fewer events). Recapitation provides other benefits as well, since it means that neutral burn-in can be deferred until after forward simulation is complete, and can even be conducted as an afterthought on existing model output. It also allows the non-neutral forward simulation to run without a burn-in history needing to be loaded (likely making it faster and leaner), and allows one to avoid the question of how many generations must be simulated for complete burn-in. It is worth noting that the coalescent (and thus recapitation) does not produce identical results to forward simulation of a Wright–Fisher model, but the differences are small and are mostly in the pattern of the most recent branches (Wakeley et al., 2012; Bhaskar et al., 2014); using recapitation as an approximation for neutral forward simulation should therefore produce practically identical results as long as the forward portion of the simulation runs for at least a few generations. Similarly, although spatial models differ substantially from the standard coalescent, this difference is mostly seen in the more recent portion of the trees; lineages that have “mixed” across the species range without coalescing behave statistically like lineages in a randomly mating population (Wilkins, 2004; Matsen and Wakeley, 2006). Recapitation with an unstructured coalescent should therefore be a good approximation to pre-existing diversity in a spatial simulation as well.

Note that constructing a burn-in history with recapitation is only equivalent to a period of forward simulation if the burn-in period is completely neutral. If a non-neutral burn-in to equilibrium is needed, the best approach is probably to run the burn-in period in SLiM with tree-sequence recording turned on and neutral mutations turned off (thus avoiding the cost of simulating the neutral mutations during burn-in, as in Example 1). If a neutral burn-in is desired, but the neutral mutations are then needed by the non-neutral portion of the simulation (perhaps because some of the neutral mutations become non-neutral due to an environmental change), one might simulate the burn-in period with the coalescent in msprime (including mutation), and then save the result as a `.trees` file using pyslim; one could then read that `.trees` file into SLiM to provide the initial state for further simulation. These techniques go beyond what we have space to illustrate here, but the manual for SLiM 3 provides further recipes showing the use of tree-sequence recording. Since it is possible to move simulation data with full ancestry records back and forth between msprime and SLiM, one can imagine many ways to combine the two to leverage their strengths while avoiding their weaknesses.

## Discussion

We have integrated support for tree-sequence recording (Kelleher et al., 2018) into the popular SLiM forward simulation software package. Tree-sequence recording can now be enabled in any SLiM simulation, and the results output to a `.trees` file that can be loaded into Python for further simulation or analysis using the msprime package (a part of the tskit framework). We have also extended the tree-sequence recording method to allow the recording and output of mutations that arise during forward simulation.

We provided four examples demonstrating the power of the tree-sequence recording method. The first example, of a simple neutral model, showed how to enable tree-sequence recording with a few trivial modifications to a SLiM model’s script. The second example illustrated the use of recorded tree sequences in post-simulation analysis in Python to estimate the characteristic reduction in neutral diversity expected around functional regions due to background selection. The third example mapped the mean true local ancestry along the chromosome in a model of the admixture of two subpopulations, again using post-simulation Python analysis. Finally, our fourth example illustrated the use of msprime to “recapitate” a SLiM run, using the coalescent to construct a neutral burn-in period after the completion of forward simulation.

All of these examples illustrated the large performance benefits that can be achieved with tree-sequence recording. Indeed, for very large neutral simulations our timing comparison indicated that the speedup due to tree-sequence recording can exceed two orders of magnitude, and can put the performance of forward simulation on par with an efficient coalescent-based simulation such as msprime (Example 1). For a large simulation with many non-neutral mutations, we still observed a speedup of more than an order of magnitude (Example 2); simulations with a lower density of non-neutral mutations should benefit even more. Similarly, compared to standard forward simulation methods, using recapitation to construct a neutral burn-in period provided a speedup of more than five orders of magnitude (Example 4), and using the tree sequence to obtain true local ancestry information provided a speedup of more than six orders of magnitude (Example 3). Memory savings observed in these models were also quite substantial.

Although we have not made use of it in these examples, SLiM records substantial metadata in the tree sequence it outputs about genomes, individuals, and mutations. This includes sex, age, and spatial location of remembered individuals, and times of origination and selection coefficients of mutations. This information, along with the tree sequence, could enable substantially more detailed dissection of evolutionary trajectories. Access to this SLiM metadata is mediated by the new pyslim package that bridges SLiM and msprime. Furthermore, the `.trees` file contains all of the information necessary to reconstruct the internal state of the simulation in SLiM, so it can be loaded back into SLiM, examined graphically using SLiMgui, and even used as a starting point for further simulation (with some caveats; see the manual).

Tree-sequence recording is not a panacea. Models that do not involve neutral mutations will not realize a speed benefit from tree-sequence recording’s ability to defer neutral mutation overlay; in fact, they will run more slowly, since the overhead of recording will not be compensated by eliminating neutral mutation simulation. Models that involve a very high recombination rate relative to the mutation rate may also not see a speed benefit from tree-sequence recording, since tracking the recombination breakpoints can become very time-consuming; informal tests indicate that this becomes important, for neutral simulations, when the recombination rate is two or more orders of magnitude larger than the mutation rate, however, so it may not be a practical concern for most models. Even if simulation performance is not improved by tree-sequence recording, the ancestry information provided by the tree sequence may still speed up analysis or provide additional statistical power, which can also be quite important in reducing total runtimes. The benefit of tree-sequence recording also depends upon factors such as the proportion of neutral to non-neutral mutations, the distribution of fitness effects from which the non-neutral mutations are drawn, the genetic architecture, the frequency with which tree-sequence simplification is performed, and many other factors. In practice, it may be worthwhile to simply compare the performance of both methods for a particular model; it is difficult to distill any reliable rule of thumb. These considerations were discussed further in Kelleher et al. (2018).

A commonly used technique for speeding up large forward simulations is model rescaling. This involves scaling down the population size (*N*) by some factor *Q*, while scaling up the mutation rate (*μ*), the recombination rate (*r*), and selection coefficients (*s*) by the same factor; this holds many common population-genetic parameters constant, since they involve products of these variables (e.g., *Nμ*, *Nr*, and *Ns*). Since these factors (as well as genetic drift) are *rates*, one generation in the rescaled model corresponds to *Q* generations in the original model. Therefore, rescaling by a factor *Q* can provide a speedup of up to a factor of *Q*^2^ due to the *Q*-times smaller population size *and* the *Q*-times smaller number of generations that need to be simulated. However, this technique has important limitations, because it can introduce artifacts due to the discretization of mutation frequencies and of time. For example, if a model with an original population size of *N* = 10,000 were rescaled to a model with *N* = 100, the smallest possible mutation frequency will also have increased from 0.00005 to 0.005, which could severely affect studies in which one is concerned about the behavior of low-frequency polymorphisms. There are more serious issues related to the process of adaptation; since rescaled values of *s* are larger, rescaling has the net effect of substituting many mutations of small effect with a single one of large effect (with *Q=*100, replacing 100 mutations with *s*=0.001 by a single one of *s=*0.1). Thus, rescaling must not be taken too far, and careful comparisons are needed between the unscaled and the rescaled model to ensure that results are not altered by rescaling artifacts. The SLiM manual (Haller and Messer, 2016) has an extended discussion of model rescaling and provides instructive examples. Since tree-sequence recording does not introduce such artifacts, it probably ought to be used to full advantage before any model rescaling is applied. If that does not bring the desired simulation within practical computational bounds, rescaling may be used in conjunction with tree-sequence recording, but with the same caveats mentioned above. Note, however, that the effectiveness of combining both strategies is hard to predict, since the increased recombination rate in the scaled model means that roughly the *same* number of recombination events must be recorded.

Although tree-sequence recording is not appropriate in every model, the examples we have presented demonstrate that the performance gains it provides can make simulations possible that would previously have been beyond reach, opening up new horizons for exploration. The software packages used here – SLiM, msprime, Python, R – are all free and open-source, and the examples and analyses shown here are all available on GitHub. We hope that the practical examples we have provided will raise the level of awareness among evolutionary biologists regarding this exciting new method.

## Acknowledgements

Thanks to Kevin Thornton and Dom Nelson for helpful discussions. This work was supported by funding from the College of Agriculture and Life Sciences at Cornell University, Predator Free 2050 (grant SS/05/01), and the NIH (grants R21AI130635 and R01GM127418) to PWM; by funding from the Sloan Foundation and the NSF (under DBI-1262645) to PLR; and by the Wellcome Trust (grant 100956/Z/13/Z) to Gil McVean for JK.

## Data Accessibility

SLiM 3 is available online at https://messerlab.org/slim/. It is open source, under the GPL 3.0 license, and its source code is on GitHub at https://github.com/MesserLab/SLiM. msprime is available online at https://pypi.org/project/msprime/. It is open source, under the GPL 3.0 license, and its source code is on GitHub at https://github.com/tskit-dev/msprime. pyslim is open source, under the MIT license, and is available on GitHub at https://github.com/tskit-dev/pyslim.

The examples and performance assessments presented in this paper are available on GitHub at https://github.com/bhaller/SLiMTreeSeqPub.

## Author Contributions

*We have used the CRediT taxonomy for contributions (https://casrai.org/credit/).*

BCH: Conceptualization, Investigation, Methodology, Software, Validation, Visualization, Writing – Original Draft Preparation, Writing – Review & Editing.

JG: Conceptualization, Methodology, Software, Writing – Review & Editing.

JK: Conceptualization, Methodology, Software, Validation, Visualization, Writing – Review & Editing.

PWM: Conceptualization, Funding Acquisition, Supervision, Writing – Review & Editing.

PLR: Conceptualization, Funding Acquisition, Methodology, Software, Supervision, Validation, Writing – Review & Editing.

